# Resolving the Foliar Calcium Mobility Paradox: Enhancing Foliar Calcium Transport in Tomato Using Osmotic Regulators

**DOI:** 10.1101/2025.07.02.662739

**Authors:** Eduardo Santos, Gabriel Sgarbiero Montanha, Higor José F. A. da Silva, Nicolas Gustavo da Cruz da Silva, Felipe Sousa Franco, Julia Rossato Brandão, Karina Angela Chimbo Huatatoca, Alex Virgilio, Chithra Karunakaran, Karen Tanino, José Lavres, Hudson Wallace Pereira de Carvalho

## Abstract

**Highlight:**

– *In vivo* XRF measurements in the petiole confirmed Sr transport to distal tomato tissues.
– XRF imaging revealed the distribution of Sr in tomato fruits following foliar application.
– Foliar formulations supplemented with osmotic regulators improved Sr absorption and translocation.
– Sucrose enhanced both short- and long-distance transport of Sr to tomato fruits.
– Sucrose significantly enhanced Sr²⁺ accumulation in seeds and the apical mesocarp, the regions most affected by Ca deficiency symptoms The study demonstrates a novel approach for enhancing phloem transport to fruit of foliar-applied Ca

Calcium (Ca) deficiency can impair fruit development even under optimal soil Ca levels due to its transpiration-dependent transport. Reduced fruit transpiration may limit Ca delivery to fruits, leading to lower Ca content in the fruit and diminished quality. Foliar application of Ca offers a potential strategy to mitigate these effects; however, its low mobility in the phloem often limits treatment efficacy. To address this, we employed X-ray fluorescence spectroscopy (XRF) to investigate the penetration and transport of foliar-applied Ca, using strontium (Sr) as a physiological tracer. Additionally, we evaluated the influence of osmotic regulators, sucrose, mannitol, glycerol, and potassium, on Ca transport. Results showed that Sr was effectively translocated to distal tissues. While potassium and mannitol had no significant impact on transport kinetics, sucrose and glycerol enhanced Sr movement. XRF imaging of leaf tissue revealed that Sr was primarily transported through the apoplast toward the leaf margin. Moreover, foliar application of Sr combined with sucrose significantly increased Sr accumulation in seeds and in the apical portion of tomato fruits. These findings suggest that, contrary to the common assumption of limited foliar Ca mobility, sucrose, can acts as an effective osmotic regulator, enhancing both short-range movement within leaf tissue and long-distance translocation to fruit.

## 1. Introduction

Calcium (Ca) is a macronutrient that plays a crucial role in plant metabolism, including the maintenance and stabilization of the cell wall and membrane integrity, as well as acting in signalling pathways involved in the response to abiotic and biotic stresses^1^. Nevertheless, Ca deficiency is prevalent in many crops, notably fruit species, which primarily stems from the limited mobility of Ca in the phloem and its restricted redistribution from mature tissues to actively growing sites. Since the transport of Ca within plants occurs predominantly through the xylem, and hence, is primarily driven by transpiration via the xylem, its distribution is highly dependent on water movement and environmental conditions^1–4^.

Localized Ca deficiency is often observed in developing tissues such as fruits, even in soils with adequate Ca availability. This phenomenon is attributed to the competition for Ca between developing fruits and leaves, coupled with low Ca redistribution, low mobility in phloem and the high demand for Ca²⁺ in rapidly expanding sink tissues such as fruits^3,5–7^. Calcium deficiency can result in severe physiological disorders such as blossom-end rot (BER) in tomatoes and bitter pit in apples, as well as reduced post-harvest shelf life and increased vulnerability to pathogen attack ^8–11^. These issues contribute to substantial economic losses in agricultural systems.

To mitigate Ca deficiency, foliar application of Ca has become a widely used strategy^12^. However, its efficacy is often limited, with less than 5% of applied Ca²⁺ typically reaching fruit tissues^13^. The majority of the Ca remains sequestered in shoot tissues due to its limited mobility in the phloem and strong interactions with cellular components such as pectins and phospholipids^13,14^. Since mechanisms underlying Ca uptake via foliar application and its subsequent redistribution within the plant remain poorly understood, addressing these knowledge gaps is crucial for enhancing the efficiency of foliar Ca application, which holds promise for improving crop quality and minimizing economic losses. Overcoming limitations of Ca immobility in plants needs innovative strategies to enhance its transport, particularly through improved phloem mobility.

*Solanum lycopersicum* is susceptible to BER, a physiological disorder primarily correlated with low Ca concentration in fruit, leading to tissue necrosis and reduced fruit quality ^12,15,16^. To investigate the potential of osmotic regulators, such as sucrose, potassium, glycerol, and mannitol, to facilitate Ca transport within the phloem to tomato fruit, the Micro Tom cultivar was selected as a plant model due to its compact size, short life cycle, and ease of cultivation under laboratory conditions.

Although Ca is traditionally considered a poorly mobile element in the phloem, we hypothesize that its mobility can be enhanced through the addition of performance additives in foliar formulations, such as osmotic regulators. These compounds are believed to increase the osmotic pressure within the phloem, thereby driving the pressure-flow mechanism and facilitating more efficient Ca transport to growing sink tissues. By establishing a steeper osmotic gradient, this approach aims to overcome physiological barriers to Ca mobility and improve its distribution and utilization. This strategy addresses critical knowledge gaps in nutrient transport dynamics and may contribute to more effective and targeted calcium nutrition in crops.^1,14,17^.

## 2. Material and Methods

### 2.1 Assessing ionic transport in petioles under vivo conditions

The effect of osmotic regulators on Ca transport in tomato plants (*Solanum lycopersicum* var. Micro-Tom) was tested in plants cultivated in a growth chamber. The plants were grown in substrate and irrigated with Modified Cakmak nutrient solution^18^ (Table S1). The environmental conditions included a 13-hour photoperiod provided by 6500 K LED lamp illumination delivering a flux of 250 µmol m^−2^ s^−1^ photons, a temperature of 25 ± 3 C, and relative humidity of 70 ± 10%. At the flowering stage (45 days old after emergence), the foliar fertilizer treatments were applied targeting the three distal leaves.

The treatment consisted of foliar applications containing CaCl_2_ 2H₂O (Química Moderna, BR) and SrCl₂·6H₂O (Êxodo Científica, BR) at concentrations of 0.1 M Ca and 0.025 M Sr, respectively, either applied individually or in combination with four osmotic regulators: glycerol (Rioquímica, BR), mannitol (Synth, BR), KCl (J.T. Baker), BR), and sucrose (Merck, DE), each at 13 mM. These concentrations were selected based on a literature review of Ca foliar application studies (not shown), which indicated an average foliar solution concentration of 0.125 M Ca. In this study, 20% of the Ca was replaced with Sr, employed as a physiological analog for Ca. The negative control was treated with water. Detailed concentrations and dosages of each foliar solution are provided in Table 1. Fertilizers were applied twice, with a 74-hour interval between applications, using a mini airbrush (Fig. S1A). Each application delivered 1 mL of solution per plant, totaling 2 mL per plant over the treatment period.

**Table 1.** Concentration and dose of foliar solution used *in vivo* Sr foliar transport experiment. In each foliar formulation was added 1% (v/v) mineral oil adjuvant.

| Treatment | Concentration (g L<sup>-1</sup>) |  |  | Dose (mg plant<sup>-1</sup>) |  |  |
| --- | --- | --- | --- | --- | --- | --- |
|  | Ca | Sr | Osmotic regulator | Ca | Sr | Osmotic regulator |
| Control- Water | 0 | 0 | 0 | 0 | 0 | 0 |
| SrCl<sub>2</sub> |  |  | 0 |  |  | 0 |
| SrCl<sub>2</sub> + glycerol |  |  | 0.82 |  |  | 1.64 |
| SrCl<sub>2</sub> + Mannitol | 4 | 2.2 | 1.63 | 8 | 4.4 | 3.26 |
| SrCl<sub>2</sub> + Potassium |  |  | 1.3 |  |  | 2.6 |
| SrCl<sub>2</sub> + sucrose |  |  | 3.06 |  |  | 6.12 |

The analysis was performed in the fertilized leaf’s petiole over 243-h using an *in-house* X-ray fluorescence spectrometer (Spectrometer for In Vivo Plant Analysis, SIPA, Brazil). The equipment is furnished with a silicon drift detector (Ketek, AXAS-68 D, Germany) and 4 W Ag X-ray tube (Amptek, Mini-X, USA) operating at 35 kV and 110 µA with a 25 µm thick Cu primary filter selected^19^. On each measurement, the dwell time was 120 seconds the Sr Ka peak was normalized by the Rh Ka Compton scattering. The Sr limit of detection (LOD) was calculated using the equation 8.456*√BG, where BG represents the background signal. The assay was carried out using five biological replicates.

### 2.2 Assessing short-distance foliar ionic transport

To characterize the short-distance transport of Ca using its physiological tracer, strontium (Sr)^20^, synchrotron-based XRF mapping was recorded in tomato leaves and petiole cross-sections. Tomato plants were grown in the University of Saskatchewan greenhouse (52°08’20.3“N, 106°37’56.0”W), and independent experiments were conducted to assess the leaf and petiole tissues. Therefore, for leaf samples, foliar Sr solutions at 0.1 M Sr, with or without sucrose, were applied to the adaxial surface of the leaf blade using a cardboard template to ensure uniform distribution along a defined midline (Fig. S1B). For petiole analysis, entire leaves were treated with the same Sr solutions (Fig. S1C). The leaf and petiole tissues were collected at 26 h and 36 h after the treatment application, respectively, embedded into OCT medium and cryofixed using liquid nitrogen. The resulting blocks were then cryo-sectioned at −25°C to yield 30-µm slices.

The XRF imaging was conducted at the BioXAS beamline of the Canadian Light Source (Saskatoon, Canada). The beamline was operated in micro-mode configuration, with excitation energy of 16.5 keV. The Vortex 3E detector was positioned at a 45° angle and 2 cm distance to the incident beam. The images were recorded for 150 ms point^-1^ by a 5 µm beam. Data were normalized by I_0_ (ion chamber before sample) and corrected by Compton scattering using PyMca software^21^.

### 2.3 Assessment of long-distance leaf-to-fruit ionic transport

The greenhouse-based experiment was conducted at the University of Saskatchewan (52°08’20.3“N, 106°37’56.0”W) between September 2023 and January 2024. Plants were grown under natural light supplemented with bulb lamps to maintain a 13-hour light per 11-hour dark photoperiod. A commpletely randomized design (CRD) was employed. The cultivation followed the same protocol used in the first experiment. At 45 days after sowing, plants received two foliar applications: at the flowering stage and two weeks later. Foliar treatments consisted of Sr^2+^ with or without sucrose, at concentrations of 8.76 g L⁻¹ Sr and 3.06 g L⁻¹ sucrose. These Sr concentrations were higher than those used in the initial experiment to ensure adequate Sr accumulation in fruits for synchrotron-based XRF mapping. Each spraying delivered 1 mL of solution per plant, amounting to a total Sr dose of 17.52 mg plant^-1^.

At the ripening stage, tomato fruits were harvested for synchrotron-based XRF analysis. The fruits were manually sliced and stored in the freezer at −80°C, followed by a freeze-drying procedure for structural and elemental distribution preservation. Chemical maps of Ca and Sr distribution were recorded at the BioXAS beamline of the CLS, configured in macro mode. These maps were acquired with incident energy at 16.5 keV, a pixel size of 10 µm, and a Dwell time 150 ms. Fluorescence signals were detected using the 4E Vortex detector. Data were normalized by Compton scattering and processed by PyMca^21^ and OriginLab software. Three biological replicates per treatment were used.

The remaining fruits were collected and processed for bulk Sr quantification. Samples were weighed, washed, oven-dried to constant weight, re-weighed, ground, and pressed into pellets. Sr concentrations were determined using the SIPA XRF equipment operating at 35 kV and 110 µA with a 25-µm Cu primary filter selected^19^. Spectra were collected over 120 seconds per sample. A standard addition calibration curve was employed to convert XRF intensity into Sr concentrations (mass/mass). The calibration curve is shown in Fig. S2.

## 3. Results

### 3.1 Short-distance Transport: i*n vivo* and tissue-resolved assays

The *in vivo* transport experiment evaluated the effects of osmotic regulators, i.e., mannitol, potassium (K), glycerol, and sucrose, on foliar Sr transport kinetics in tomato plants. All treatments containing Sr presented higher Sr^2+^ than the negative control (deionized water) (Fig. 1). The addition of K^+^ and mannitol did not enhance Sr transport relatively to SrCl₂ (Fig. 1A and 1C), whereas an increased Sr transport was observed following the second application of sucrose and glycerol, with the most pronounced signal enhancements observed at 75 and 96 hours after the second application for sucrose and mannitol, respectively (Fig. 1B and 1D).

**Fig. 1.**
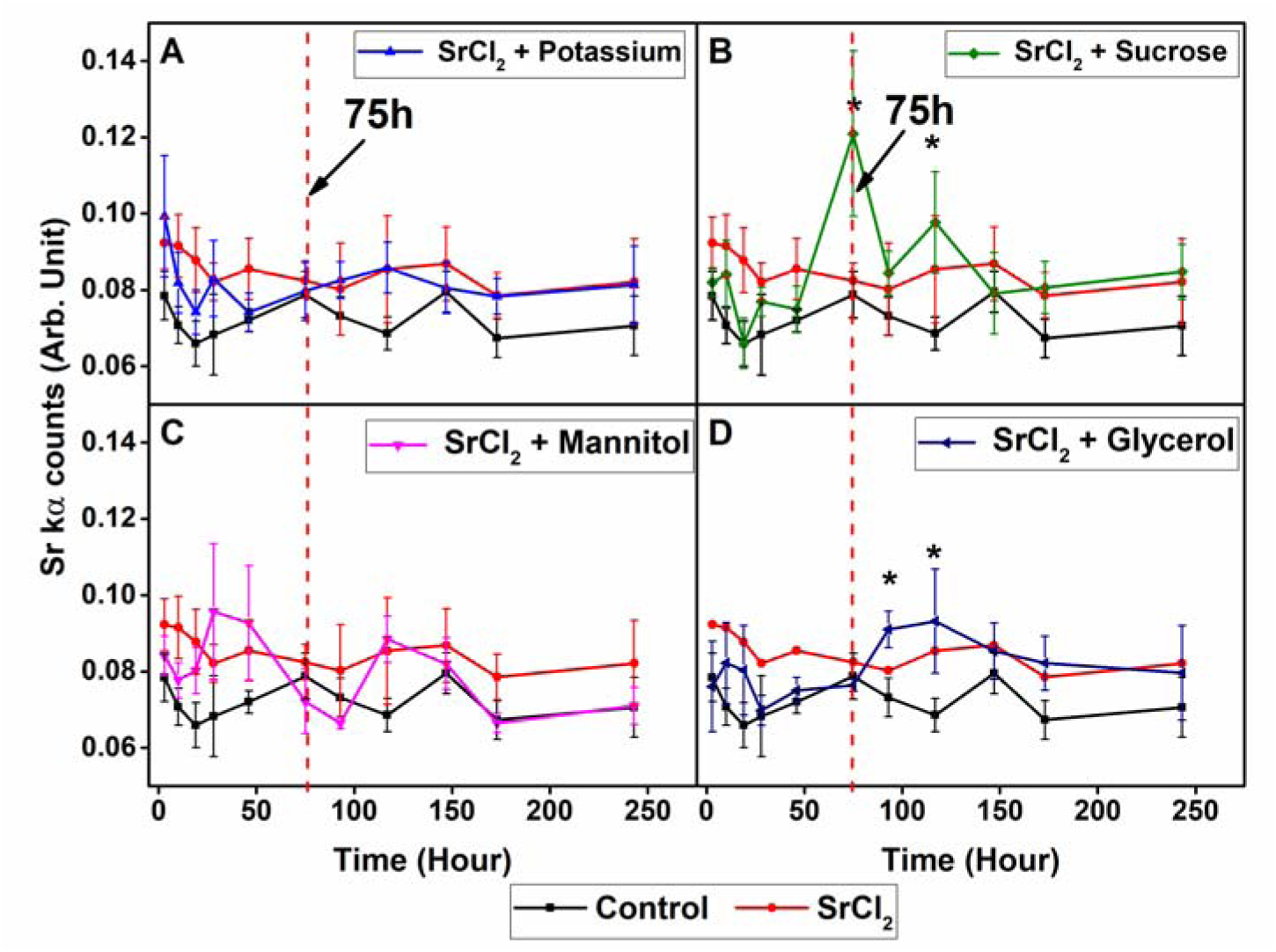
*In vivo* XRF measurement of foliar Sr transport kinetics through the petioles of tomato leaves exposed to SrCl₂ alone (represented by the red line) or combined with osmotic regulators, i.e., potassium-blue line (A), sucrose-green line (B), mannitol-pink line (C), and glycerol-navy blue line (D). The negative control consisted of deionized water and is represented by the black line. Data represents the mean ± standard error of five (n=5) independent biological replicates for all the treatments except the control, where n=4, due to stem collapse during the assay. Arrows represent the measurement after the second application, and the asterisks highlight the increased transport of Sr^2+^.

Moreover, it is important to highlight the distinct patterns observed among treatments. Those containing K^+^ and glycerol addition exhibited an early Sr peak within the first 24 hours, followed by a decline and stabilization at levels lower than the positive control (SrCl₂ alone) (Fig. 1A and 1C). Conversely, the mannitol addition showed a gradual increase of Sr transport up to 48 hours after foliar fertilization, whereas the sucrose treatment presented a pronounced increase in Sr intensity following the second application, resulting in the highest overall signal among all treatments (Fig. 1B). The negative control, which received no foliar Sr treatment, exhibited a constant Sr intensity throughout the experiment, reflecting baseline Sr uptake *via* roots due to naturally occurring Sr in the growth substrate. Figure S3A shows the cumulative signal of Sr revealing that sucrose was the most effective addive to increase Sr transport compared to the other osmotic regulator treatments (K, mannitol, and glycerol). Based on these results, sucrose was selected as the osmotic regulator for subsequent experiments.

Furthermore, the short-distance transport of Sr after foliar application was also investigated by synchrotron-based X-ray fluorescence spectroscopy mappings of leaf cross-sections from three regions: apex, center, and base, with the center corresponding to the treated area (Fig. 2). The distribution of Ca across the three regions and control was relatively uniform, as expected for a nutrient mostly found adhered to the cell wall’s middle lamella (Fig. S4 and S5). In contrast, Sr distribution varied significantly between the apex and basal regions of treated leaf, with the highest intensities observed in the apex regions (Fig. 2). In the control leaves (water treatment), Sr hotspots were observed primarily in the vascular bundles, co-localizing with Ca hotspots (Fig. S5).

**Fig. 2.**
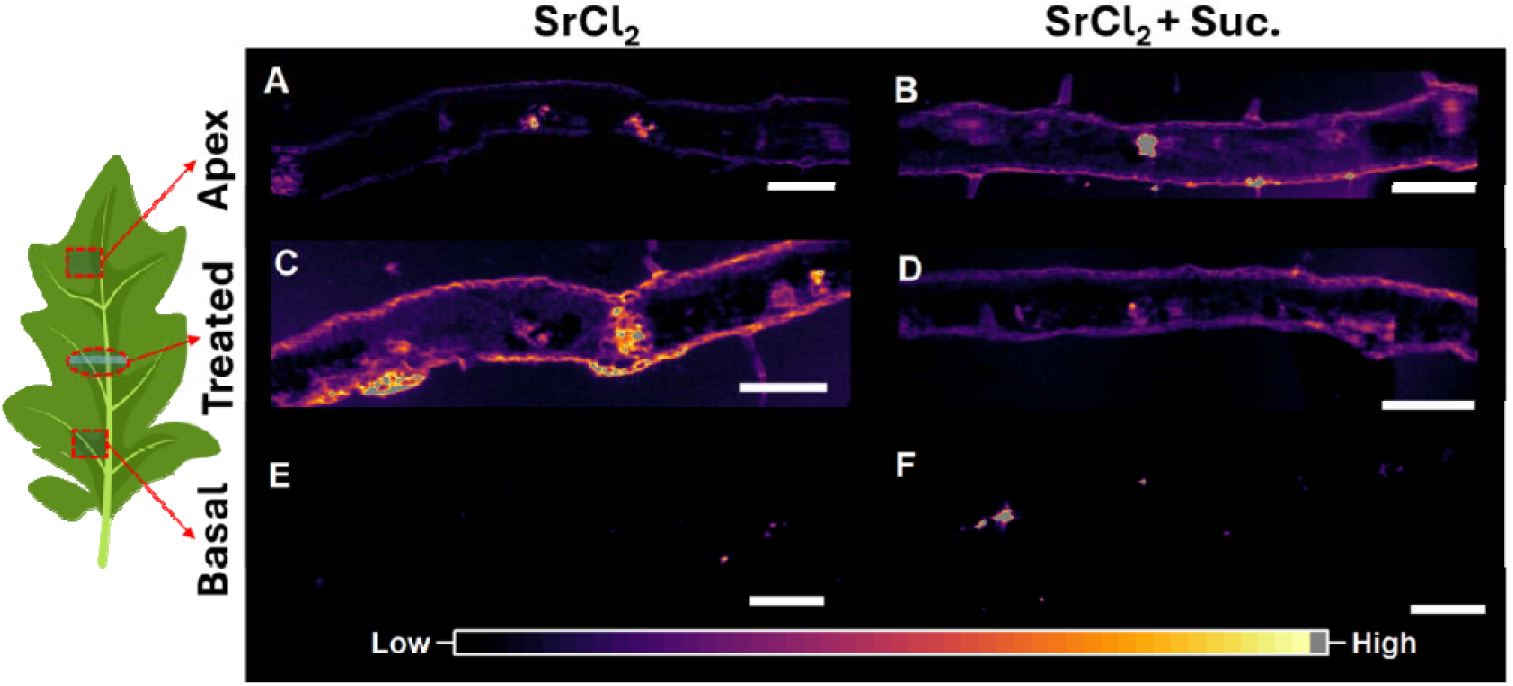
Distribution and intensity of Sr cross-sections of treated tomato leaves. Three regions of the leaf were sampled and mapped, distal and proximal from the treated area that correspond to the apex and basal regions of the leaf, respectively. The red box on the schematic leaf indicates the sampling region, with the red arrow indicating the corresponding XRF map location of the apex (A and B), treated region (C and D), and basal (E and F). The red oval indicates the treated region. The intensity scale for Sr distribution maps is consistent across all maps. The scale bars represent 250 µm.

The addition of sucrose to the foliar solution enhanced the Sr transport to the leaf apex (Fig. 2 B), as evidenced by higher Sr intensities compared to those tissues exposed to Sr alone (Fig. 2A). Conversely, no differences in Sr distribution were observed in the basal region as a function of the sucrose addition (Fig. 2 E and F). This pattern indicates transport of Sr towards the tip of the treated leaf. XRF maps of the leaf also revealed Sr hotspots that co-localized with vascular bundles and Ca hotspots. This distribution pattern was consistent in both foliar treatments and water-treated leaves, suggesting this represents Sr typical distribution in leaves after root uptake of Sr (Fig. 2 and Fig. S5). Interestingly, the XRF images of leaf cross-sections also revealed Sr signals were concentrated around cell peripheries (palisade cells), closely aligning with Ca distribution patterns in the cell walls (Fig. 2 and S4), suggesting that its transport occurred primarily *via* the apoplast Sr.

Translocation of Sr^2+^ from the site of foliar application to adjacent tissues was confirmed by the XRF maps recorded on the cross-sections of petiole tissues from leaves exposed to Sr solutions (Fig. 3). Figures 3E and F show a similar distribution pattern for Sr treatment alone and with sucrose. The Sr signals were distributed throughout the petiole, with particularly high intensities observed in the lateral vascular bundles, denoted with arrows, which are anatomically connected to the leaf edge bundles ^22,23^. Similar to the leaf cross-sections, Sr hotspots were detected in both control and treated samples. However, in Sr-treated plants, Sr was more homogeneously distributed across the petiole tissue, whereas in the control group, Sr hotspots appeared sparsely and strongly co-localized with Ca signals (Fig. 3).

**Fig. 3.**
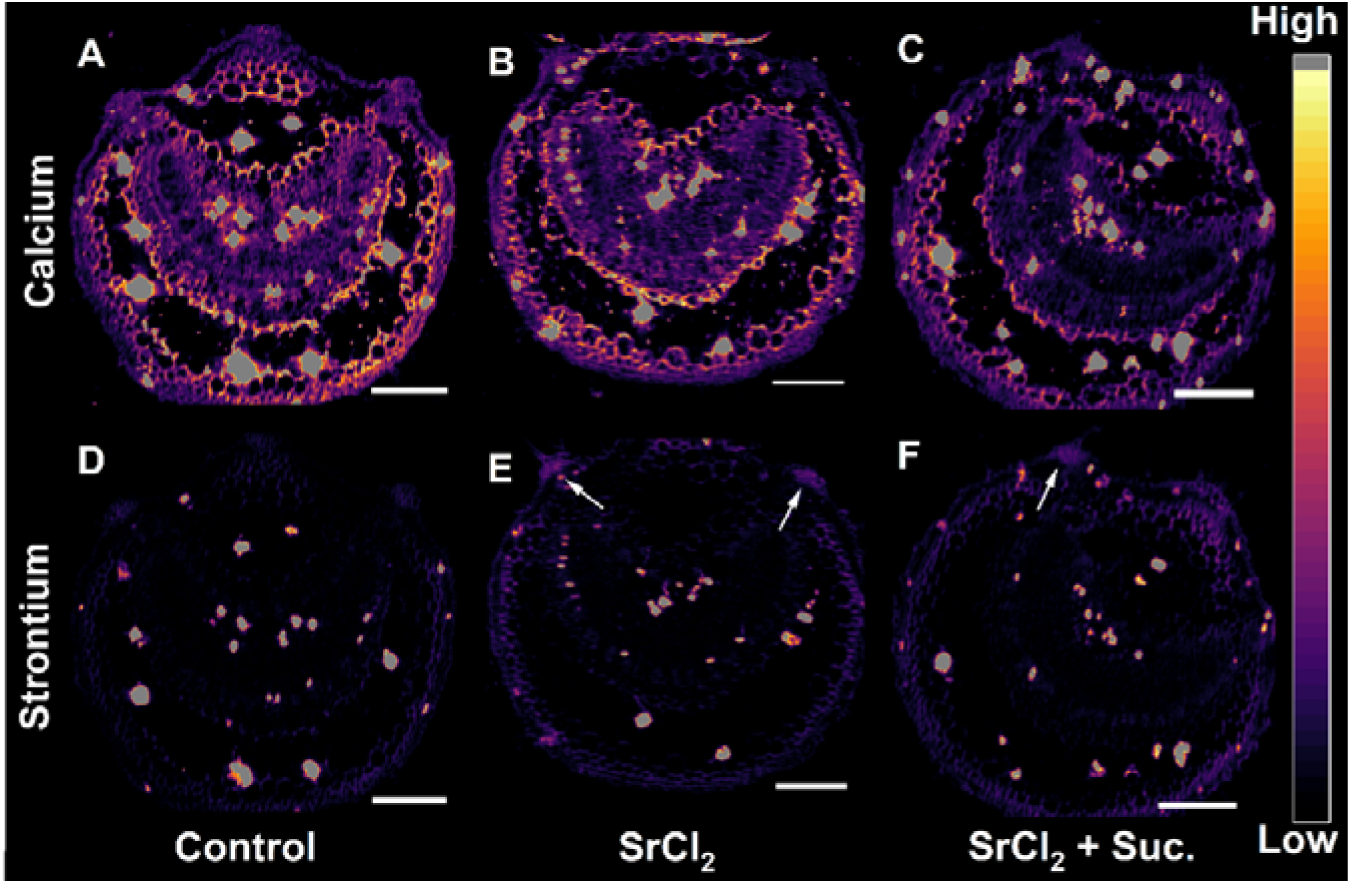
Distribution of Sr in petiole cross-sections of tomato plants treated with different foliar solutions. Control group, plants treated with water (A, D). Petioles of plants treated with SrCl₂ alone (B, E). Petioles of plants treated with SrCl₂ combined with sucrose as an osmotic regulator (C, F). The arrows indicate the lateral vascular bundle. Sr signals are represented using a color scale from blue (minimum intensity) to red (maximum intensity). Scale bars = 50 µm.

### 3.2 Long-distance transport

The impact of foliar Sr application, with and without sucrose, on tomato yield, biomass, and Sr translocation was assessed. Foliar application of Sr without sucrose led to a higher fruit yield; however, this increase was not reflected in fruit dry mass, which remained similar to both the negative control (Ctrl) and the Sr + sucrose (SrCl_2_ + Suc.) treatment (Fig. 4). All treatments that received Sr showed significantly greater Sr concentration and accumulation in the fruits compared to the control. Nonetheless, no significant differences were observed among the Sr-treated groups.

**Fig. 4.**
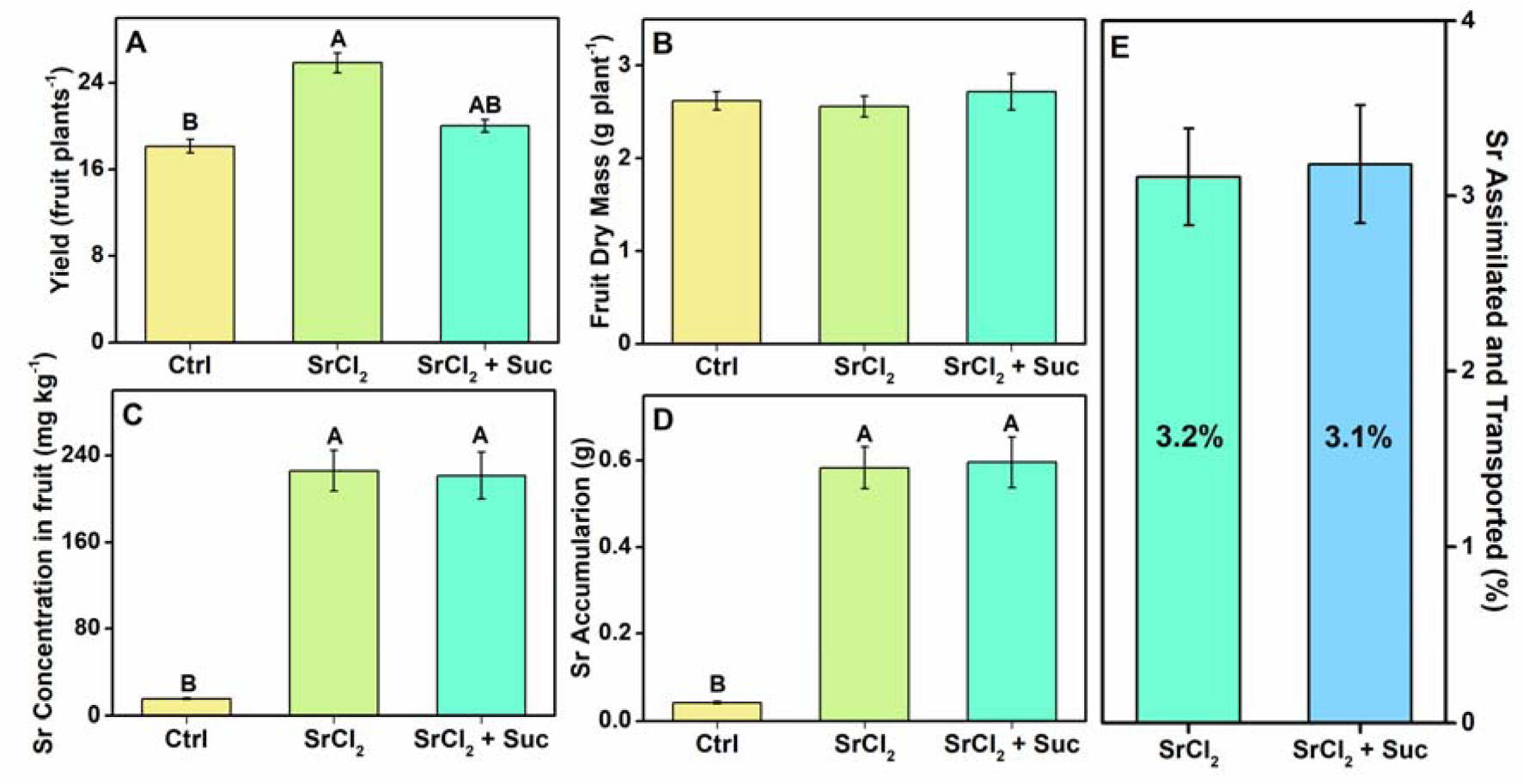
Effect of foliar Sr application with and without sucrose on tomato yield, biomass, and Sr translocation. Tomato plants were subjected to Ca restriction (5% of the standard nutrient solution Ca concentration) one week before foliar application. The treatments included a control group (Ctrl), which received only water, and two Sr treatments: Sr alone (SrCl_2_) and Sr with sucrose (Suc). The foliar solutions were applied to both leaves and fruits without fruit coverage. Evaluated parameters include fruit yield (A), dry biomass (B), Sr concentration in fruits (C), Sr accumulation in fruits (D), and Sr transport efficiency to fruits (E). Different letters indicate significant differences between treatments (p < 0.05, ANOVA followed by Tukey’s test). Each treatment was replicated in six biological samples, and error bars represent the standard error of the mean.

Moreover, the XRF maps of tomato fruits reveal similar Sr distribution patterns in the tissues subjected to foliar Sr treatments with and without sucrose (Fig. 5). In both cases, Sr was primarily concentrated in the vascular bundles and placental tissues, with a clear accumulation also in the seeds (Fig..5 B and E). However, the fruits of plants exposed to SrCl₂ + sucrose demonstrated more intense and widespread Sr signals throughout the internal tissues, particularly in the central columella and surrounding placenta, compared to those treated with SrCl₂ alone. While there was no evidence of alteration in Ca distribution in Sr treatments, the XRF maps of Ca highlight the preservation of fruit structure, the presence of calcium oxalate crystals in the central columella, indicating native Ca²⁺ deposition sites, not influenced by foliar treatments, and a signal absence of Ca in the seeds (Fig. 5 A and D). The overlap of the Ca and Sr XRF maps emphasizes the contrast in the distribution of both elements in the fruits (Fig. 5 C and F). The same distribution pattern for both elements, Ca and Sr, was observed in replicates 2 and 3 (Fig. S6 and S7).

**Fig. 5.**
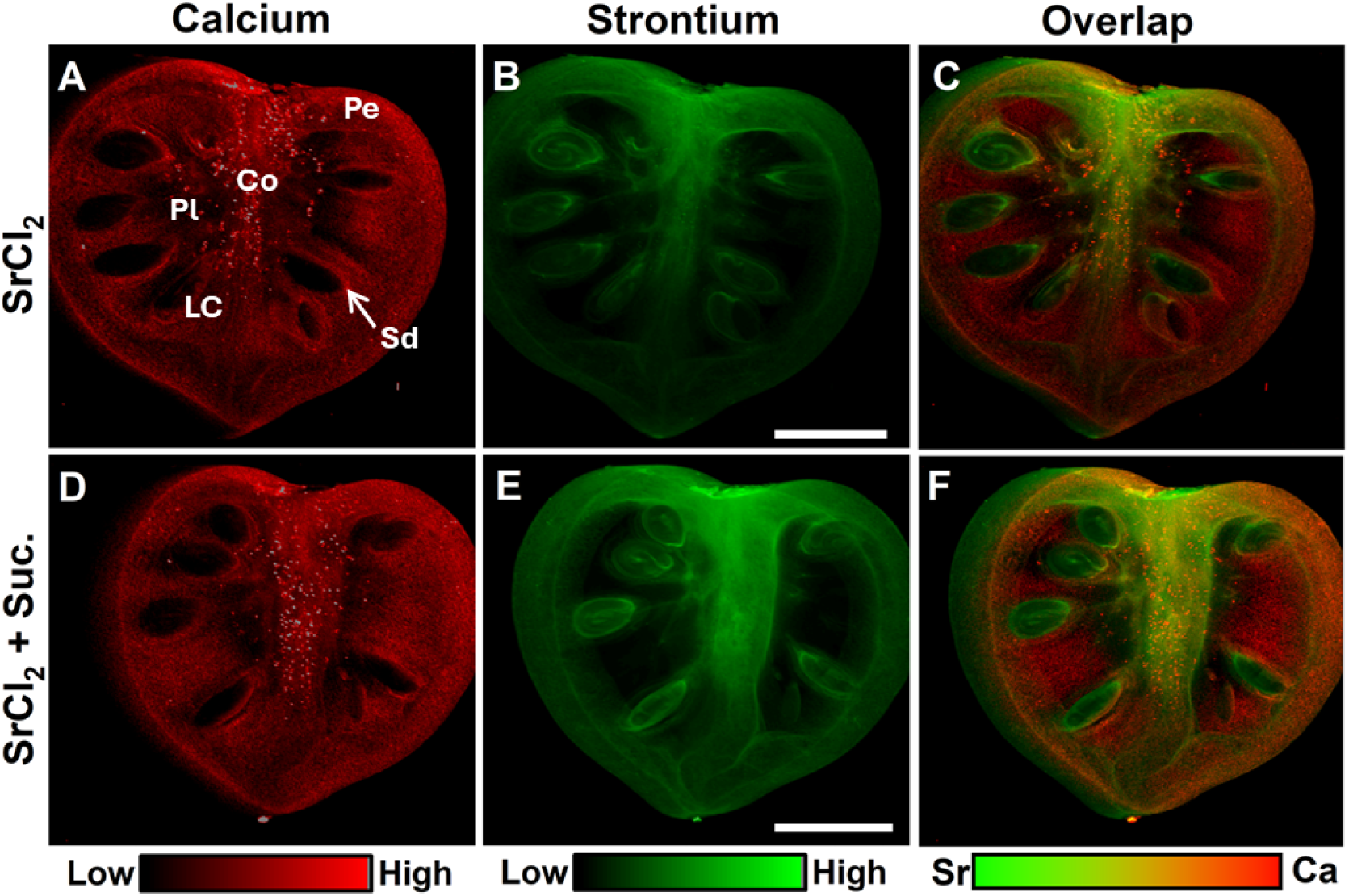
Distribution of Sr and Ca in the transverse cross-section of tomato fruits from plants subjected to foliar application of SrCl₂ alone (A-C) and with sucrose (D-F). XRF map of Sr with a green scale bar ranging from low Sr intensity (black) to high intensity (light-green) (A and D). Ca distribution map with a color scale bar ranging from black to light red, indicating low (black) and high (light red) Ca intensity, respectively. Overlap of both Ca and Sr elements (C and F), corresponding to green and red, respectively. Tomato fruits manually sectioned transversely were imaged at the BioXAS beamline of CLS. Different tissues were identified and represented as pericarp (Pe), collumela (Co), placenta (Pl), locular cavity (LC), and seeds (Sd). The fruits exhibit a more uniform Sr distribution throughout the entire fruit, with particularly higher intensities observed in the seeds, vascular bundles, and placenta. Scale bars represent 3 mm.

Despite full exposure of the fruits during foliar application, no gradient of Sr intensity was observed in the pericarp, even though such a pattern would typically indicate epidermal absorption (Fig. 5 B and E). Additionally, the control group (water-treated plants) exhibited a noisy Sr^2+^ map with a localized hotspot near the peduncle insertion at the fruit base, while Ca maps showed a consistent distribution pattern across all replicates (Fig. S8).

The semi-quantitative analysis of the XRF Sr maps reveals the average Sr signal intensity across different fruit tissues. Sr concentration followed the pattern: basal mesocarp > columella > seeds > apical mesocarp (Fig. 6). Notably, the sucrose-treated fruits exhibited higher Sr levels compared to the Sr-only treatment, particularly in the seeds (Fig. 6B) and apical region (Fig. 6C), where the differences were statistically significant at the 5% confidence level.

**Fig. 6.**
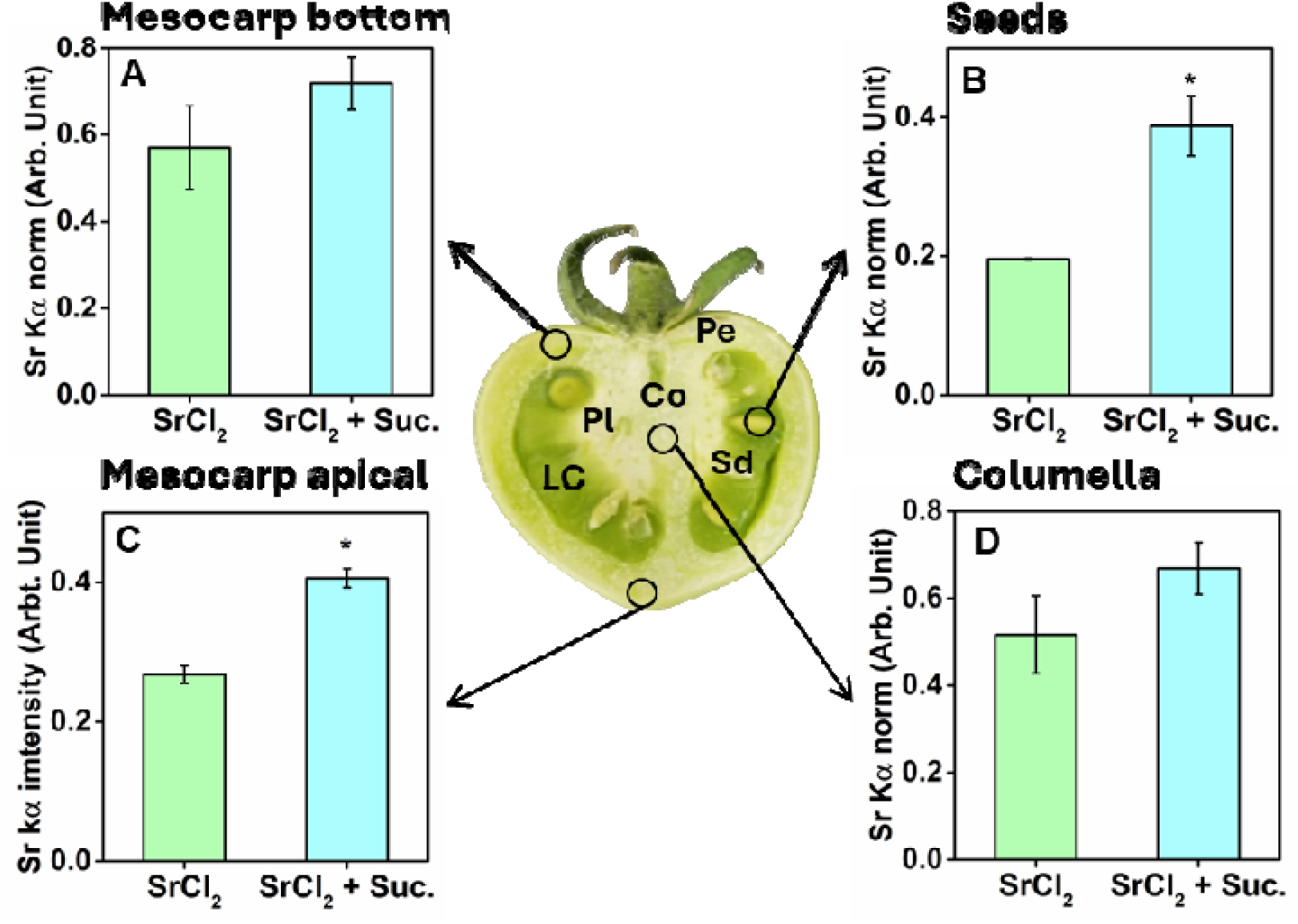
Average intensity of Sr kα across different tissues of tomato fruits from plants treated with either a foliar solution of Sr alone or with sucrose. The Sr intensity in the basal mesocarp (A), seeds (B), apical mesocarp (C), and collumela (D) was determined using X-ray fluorescence (XRF) mapping. The different tissues were identified and labeled as pericarp (Pe), collumela (Co), placenta (Pl), locular cavity (LC), and seeds (Sd). Asterisks indicate significant differences between treatments (p < 0.05, ANOVA followed by Tukey’s test). Error bars represent the standard error of the mean.

## 4. Discussion

In this study, Sr was used as a physiological tracer for Ca, providing valuable insights into the foliar absorption and transport mechanisms of mineral nutrients in tomato plants. The decision to use Sr as a tracer for Ca stems from their chemical similarities, as both elements belong to Group 2 of the periodic table and share comparable ionic radii and charge (+2)^24^. This similarity allows Sr to mimic the physiological behavior of Ca in plants, such as foliar absorption and transport ^25–27^, which facilitates tracking Ca movement under *in vivo* conditions.

Although plants can absorb Ca through the leaves^28^, their long-distance transport in the phloem is limited by Ca physiological and physicochemical characteristics ^4,29^. In plant cell walls, Ca ions interact with the carboxyl groups (-COO⁻) of pectin’s galacturonic acid residues, forming pectate, causing Ca immobilization^4,14,17,30^. This cross-linking creates a gel-like structure that provides mechanical strength, facilitates cell-to-cell adhesion, and enhances resistance to enzymatic degradation. In membranes, Ca is essential for maintaining structural integrity, mediating signal transduction, and regulating cellular transport and stress responses^1^. Its dual role as a structural component and a dynamic messenger, combined with its high reactivity, contributes to its limited mobility and low remobilization dynamics within the phloem. These characteristic underscores the challenges associated with Ca redistribution in plants.

Figure 1 illustrates the effect of osmotic regulators on Ca transport following foliar application. Sucrose and glycerol treatments exhibited an increased Sr transport after the second application. Notably, among the osmotic regulator treatments, sucrose was unique in showing a statistically significant increase in transport rate compared to the control. This behaviour may be linked to the ability of osmotic regulators to enhance phloem transport of Ca by altering osmotic gradients in line with the pressure-flow hypothesis of phloem transport^13,14,31^. According to this theory, solute movement through the phloem is driven by turgor pressure gradients created by osmotic differences between source and sink tissues. It is hypothesized that exogenous sucrose was transported to the phloem, reducing the osmotic pressure in the sieve tube, thereby enhancing water flow into the phloem by osmosis from xylem and consequently promoting greater Sr transport to other tissues. Overall, our results demonstrated that sucrose was the most effective osmotic regulator to enhance Ca transport through the phloem. Therefore, this treatment was selected for further characterization of both short- and long-distance transport.

Short-distance transport refers to the movement of water, nutrients, and other molecules across small distances within plant tissues, typically at the cellular level^31^, which occurs either through diffusion, osmosis, or active transport^31^. To investigate this phenomenon after foliar fertilization, a localized foliar application was performed on the middle section of the leaves, followed by an analysis of the sections above and below the application site (Fig. 2). Higher Sr concentrations were observed in the leaf apex compared to the leaf base, indicating a gradient consistent with directional short-distance transport of Sr towards the leaf margins. Naturally, as xylem sap reaches the leaf, Sr follows the water flow, moving towards the leaf edges, where it accumulates ^14,17^. Additionally, despite the treatment applied to the adaxial surface, Sr was detected across all tissue layers, including the abaxial epidermis, a prerequisite for systemic translocation. This observation suggests that Sr penetrated the leaf surface and diffused throughout the tissue, most likely facilitated by apoplastic transport, since the Sr signal is associated with the cell wall (Fig. 2). Similar results were observed in sunflower and soybean exposed to Zn^32,33^. The authors suggested that the bundle sheath extension which crosses the leaf may facilitate the transport of Zn from the adaxial surface to the abaxial surface^33^.

On the other hand, in this study, long-distance transport refers to the movement of Sr from the treated tissue to other regions of the plant, as this term encompasses the transport of minerals, water, and signaling molecules across different parts of the plant, typically through the vascular system^31^. Herein, considering the direction of transport in the xylem, the applied fertilizers have likely travelled through the phloem within the petiole to reach other tissues.

Our findings demonstrate that a portion of the foliar-applied Sr was transported to other tissues. The XRF imaging of the petiole cross-sections revealed high Srconcentration in the lateral vascular bundles (Fig. 3), which connect to smaller veins within the leaf blade, forming an integrated vascular network ^22,23,34^. This vascular system may provide a pathway for Ca transport following foliar application. While the precise mechanism of Ca translocation remains unclear, previous literature highlights the limited phloem mobility of Ca ^5,14^. However, studies utilizing Sr and ^45^Ca have demonstrated that a portion of foliar-applied Ca can be transported to other tissues ^13,35,36^. Thus, the results presented here proved the transport of Ca to other tissues, which looks like it should occur through the phloem.

To the best of our knowledge, this is the first attempt showing Sr distribution in fruits following foliar application. In this experiment, Sr was applied to the entire shoot, including the fruits. The higher Sr concentration observed in the seeds suggests that the Sr was likely delivered via the phloem following foliar absorption (Fig. 6). This conclusion is further supported by the limited capacity of the Sr absorbed through the fruit epidermis to move internally within the fruit^37^. The absence of Ca inside the seeds may be associated with its major abundance in the seed coat, which is highlighted surrounding the seeds in Figure 5, as evidenced in Arabidopsis seeds^38^. Additionally, considering the thin layer of seed coat compared with the whole samples, the higher intensity of fruit tissue masked the Ca signal of the seed coat.

In this study, sucrose application enhanced the translocation of Sr to tomato fruits, particularly to the seeds and mesocarp tissues in the apical region. Although the role of sucrose in promoting mineral mobility is still emerging, previous studies have demonstrated its physiological benefits when applied on leaf. For example, foliar sucrose application has been associated with improved growth and nutritional quality in pea sprouts^39^, as well as enhanced transplant quality, crop establishment, and increased tolerance to cold and darkness in tomato seedlings^40^. Furthermore, sucrose has been reported to enhance growth, physiological performance, and tolerance in tomatoes exposed to poor light quality^41,42^.

Based on these results, it can be hypothesized that foliar-applied Ca penetrates the leaf surface and is transported toward the leaf margins via the apoplast, driven by mass flow associated with transpiration (Figure 7). Then, it is loaded into smaller veins within the leaf blade and subsequently translocated to other plant tissues. Sucrose enhances Ca loading into the phloem, considering that exogenously applied sucrose is absorbed by the leaf, enters mesophyll cells through the plasma membrane, and diffuses toward the vascular bundles. There, it is exported into the apoplast and subsequently taken up into the phloem symplast via SUC/SUT transporters, which utilize H⁺ gradients to drive active transport. This process increases the osmotic pressure in the phloem, drawing water from the xylem by osmosis ^43–45^. The resulting rise in turgor pressure enhances the bulk flow of phloem sap, thereby promoting Ca transport to sink tissues, such as developing fruits (Figure 7).

**Figure 7.**
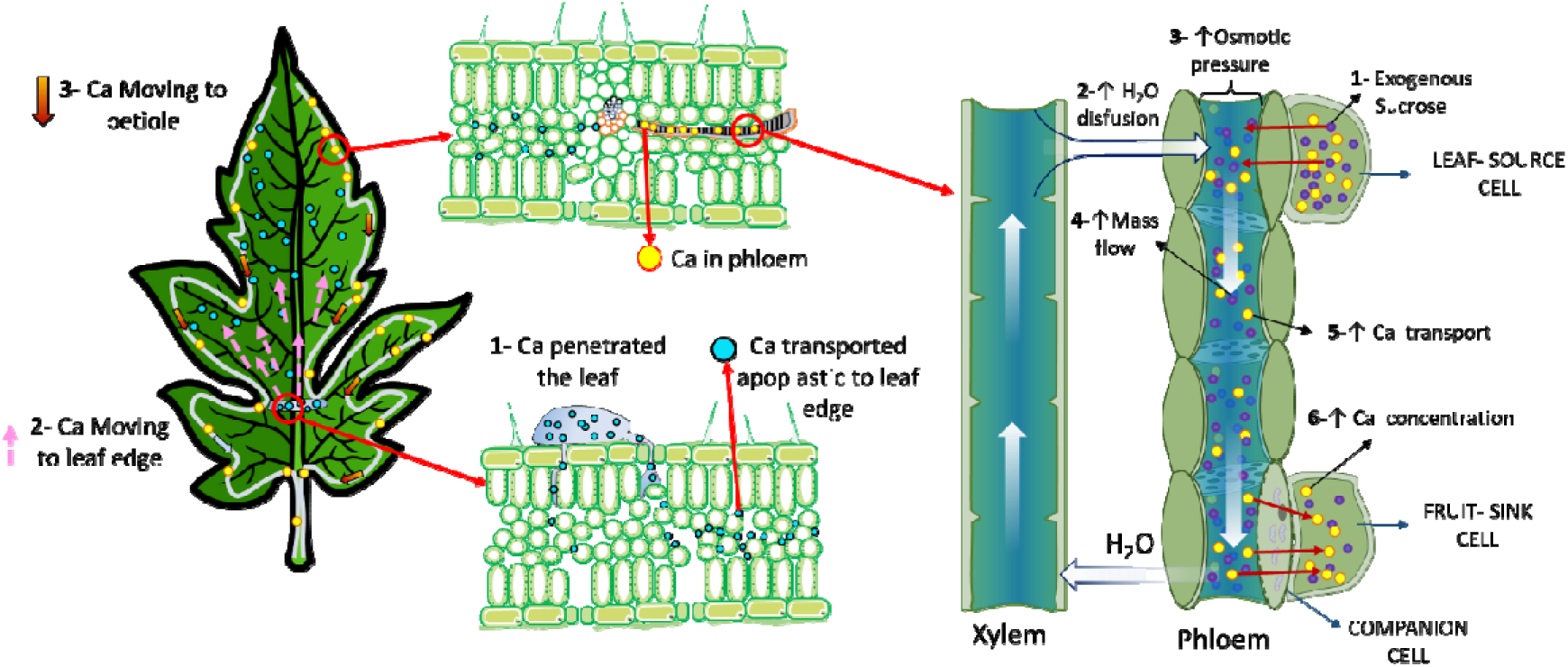
Scheme of Ca absorption and enhanced transport to fruit tissues facilitated by sucrose. After foliar application, Ca penetrates the leaf surface and crosses the cuticle and epidermal cell layers (1). Once inside the apoplast, Ca can move through the intercellular space to leaf edge following the mass flow by apoplastic transport (2). Ca is then loaded into the phloem tissues (3). The addition of sucrose, acting as an osmotic regulator, promotes water flux and may facilitate Ca movement by decreasing osmotic pressure in the phloem, thereby facilitating Ca loading and transport. As a result, Ca translocation from source leaves to sink organs, such as developing fruits, is enhanced, contributing to improved fruit quality and a reduction in calcium-related physiological disorders.

The use of sucrose to facilitate mineral transport is a novel approach. This novel strategy may increase calcium concentrations in fruits, improving the efficiency of foliar Ca application and significantly reducing crop losses caused by Ca deficiencies. Ultimately, these findings could contribute to more sustainable agricultural practices and create new oportunities for optimizing nutrient management in modern, high-yielding cropping systems.

In conclusion, this work findings confirm Sr serves as an effective physiological tracer for Ca, indicating a small but significant portion of foliar-applied Ca is transported to other tissues through *in vivo* experiments. The short-distance transport of Ca primarily occurs toward the leaf edge through the apoplast and mass flow. The addition of sucrose enhanced its movement toward the apical region. Moreover, the distribution patterns suggest Ca transport through the leaf occurred primarily *via* the apoplastic pathway. Additionally, XRF map of the petiole suggests that Ca translocation within the petiole occurs mainly through accessory (lateral) vascular bundles connected to the leaf edge veins. Our results reveal that sucrose, when used as an osmotic regulator, can significantly enhance both short- and long-distance transport of Ca, suggesting a promising approach to improving Ca mobility in the phloem.

## Supplementary data

The supplementary data include:

- **Table S1** – Nutrient solution composition used for plant irrigation.
- **Figure S1** – Experimental design for in vivo analysis and synchrotron-based XRF maps of leaf and petiole cross-sections.
- **Figure S2** – Strontium (Sr) calibration curve.
- **Figure S3** – Semiquantitative results from the in vivo analysis.
- **Figure S4** – XRF calcium (Ca) distribution maps in leaves.
- **Figure S5** – Control XRF maps of leaves.
- **Figure S6** – XRF maps of fruits from replicates 2 and 3 exposed to SrCl₂ + sucrose.
- **Figure S7** – Calcium and strontium XRF maps of fruits from replicates 2 and 3 exposed to SrCl₂.
- **Figure S8** – XRF maps of fruits from control treatments.

## Acknowledgements

We thank the BioXAS beamline of the Canadian Light Source (CLS) in Saskatoon, Canada, where synchrotron based XRF measurement were performed [proposal#39G13910] and [proposal #37G13018]. Our results reveal that sucrose, when used as an osmotic regulator, can significantly enhance to our local contact, Dr. Viorica F.Bondici for the support with measurement. We thank Prof. Dr. Felipe Klein Ricachenevsky of Departamento de Botânica, Instituto de Biociências, Universidade Federal do Rio Grande do Sul, Porto Alegre, RS. Brazil. for his valuable contribution during the revision of this manuscript.

## Author contributions

ES, CK, KT, JL, and HWPC: conceptualization; ES,GSM, HJFAS, NGCS, FSF, JRS, and KACH: investigation, AV: methodology; AV, CK, KT, JL, and HWPC; ES, CK, KT, JL, and HWPC: supervision; ES: Writing – Original Draft Preparation; ES, GSM, FSF, AV, KT, KT, JL, and HWPC: Writing – Review & Editing.

